# High-fidelity (repeat) consensus sequences from short reads using combined read clustering and assembly

**DOI:** 10.1101/2023.10.26.564123

**Authors:** Ludwig Mann, Kristin Balasch, Nicola Schmidt, Tony Heitkam

## Abstract

**Background:** Despite the many cheap and fast ways to generate genomic data, good and exact genome assembly is still a problem, with especially the repeats being vastly underrepresented and often misassembled. As short reads in low coverage are already sufficient to represent the repeat landscape of any given genome, many read cluster algorithms were brought forward that provide repeat identification and classification. But how can trustworthy, reliable and representative full-length repeat consensuses be derived from unassembled genomes?

**Results:** Here, we combine methods from repeat identification and genome assembly to derive these robust consensuses. We test several use cases, such as (1) consensus building from clustered short reads of non-model genomes, (2) from genome-wide amplification setups, and (3) specific repeat-centred questions, such as the linked vs. unlinked arrangement of ribosomal genes. In all our use-cases, the derived consensuses are robust and representative. To evaluate overall performance, we compare our high-fidelity repeat consensuses to RepeatExplorer2-derived contigs and check, if they represent real transposable elements as found in long reads. Our results demonstrate that it is possible to generate useful, reliable and trustworthy consensuses from short reads by a combination from read cluster and genome assembly methods in an automatable way.

**Conclusion:** We anticipate that our workflow opens the way towards more efficient and less manual repeat characterization and annotation, benefitting all genome studies, but especially those of non-model organisms.

## Background

In the last ten years, the amount of publicly available read data has been growing exponentially from Terabytes in 2012 to over 50 Petabytes in 2023 alone, as published in the European Nucleotide Archive (ENA statistics, accessed 08/2023). This aptly reflects the recent advances in sequencing techniques. Especially the costs of short read sequencing methods are dropping drastically and long read technologies are advancing and getting more and more accurate. This almost results in an oversupply of sequencing data and the needs for automated or at least semi-automated ways to analyse the amount of data are constantly rising. The gold standard of genomics has become to assemble a reference genome sequence; however, this can be very challenging and costly depending on the organism of interest. For species with large and complex genomes, huge amounts of data are necessary, including long reads, HiC sequencing reads or similar [1–3]. In addition to the high costs, very capable computational resources are required. Hence, especially prior to full genome sequencing projects, it is helpful to gain insight into the complexity of the target species genome in advance. One major aspect of the complexity of a genome is the fraction of repetitive DNA, which is also a main genome size determining factor [4]. Even some of the latest telomere-to-telomere genome assemblies lack the complete resolving of repetitive DNA. In contrast to full genome assemblies, to gain an overview of the repetitive DNA of a genome, only short reads and genome skimming are needed [5, 6].

Current research on repetitive DNA highlights its roles in providing genome structure, regulating transcription and pushing evolution [7–11]. One widely applied tool to identify, classify and, to a certain extent, quantify repetitive DNA from genome skimming data is the RepeatExplorer2 pipeline (RE2) [12]. Although it produces an excellent overview of the repetitive DNA fraction, however, building trustworthy repeat consensuses from its output using short reads is still a very manual, time-consuming and not reproducible process, relying mainly on the experience of the user. The RE2 output includes highly informative cluster graphs, but rather fragmented contigs of different repeat families such as transposable elements. Thus, building a comprehensive repeat consensus database with high-fidelity consensuses spanning complete or nearly complete transposable elements would be very beneficial to reduce the workload for manual curation.

However, repeat consensus reconstruction is not only needed after whole genome shotgun sequencing. It can also help to identify enriched DNA in genome-wide amplification setups. Here, we further explore our consensus building pipelines to identify and reconstruct extrachromosomal circular DNAs (eccDNAs). EccDNAs are ring-like DNAs that are physically separated from the chromosomes and have received much interest in recent years. There is still only little known about their function, but eccDNAs have been assumed to be related to aging, cancer and transposable element activation [14–17]. Here, we explore consensus-building in eccDNA circle reconstruction, building on our eccDNA identification pipeline, the ECCsplorer [13]. Similar to the default usage of RE2, the resulting ECCsplorer contigs represent rather fragmented than complete circular sequences. Therefore, reconstructing a consensus would improve the analysis of eccDNAs, since the ECCsplorer pipeline is the only published method for the *de novo* identification of eccDNAs from only short reads so far. To overcome the fragmentation of contigs, we here combine repeat clustering by RE2/ECCsplorer with assembly tools and visualization to derive high-fidelity repeat/eccDNA consensuses.

Furthermore, to complement the combination of tools, we investigate the usage of genome skimming data with only assembly tools to analyse structural features of repetitive DNA, such as linkage and separation of the highly abundant rDNA. Typically, the tandem repeats of 5S rDNA and 35S rDNA are organized in separate arrays, often on different chromosomes. In some species, however, the 5S rDNA gene is integrated into the 35S rDNA spacers, forming a so-called linked arrangement [18].

To demonstrate all mentioned strategies, we set up three different use cases. First, we use assembly tools on the default RE2 output to build comprehensive repeat consensus sequences from *Beta corolliflora*. Second, we reconstruct mitochondrial mini-circles from the *Beta vulgaris* ssp. *vulgaris* (*Beta vulgaris*) genome derived from a previous ECCsplorer output with enriched sequencing data (eccDNA-Seq). Last, we investigate the presence or absence of rDNA linkage in the two Asteraceae species *Artemisia annua* and *Tragopogon porrifolius*. For all of these use cases there is public data available. We examine all three use cases and assess the performance of the suggested workflow. Based on this, we conclude that our explorative analysis is a useful tool to semi-automatize the analysis of repetitive DNA in non-model genomes.

## Methods and Implementation

The presented repeat assembly workflow uses clustering and assembly tools to create informed consensus sequences from repeats to answer a wide variety of questions. We use the term “informed consensus” to suggest that the derived sequences are not mere averages or sequence profiles, but that they have been carefully constructed using relevant data and analysis. The workflow can be used to generate repeat databases with comprehensive consensus sequences, reconstruct eccDNA sequences from eccDNA-Seq or provide insights into structural features of highly abundant repeats (Fig. 1).

**Fig. 1:**
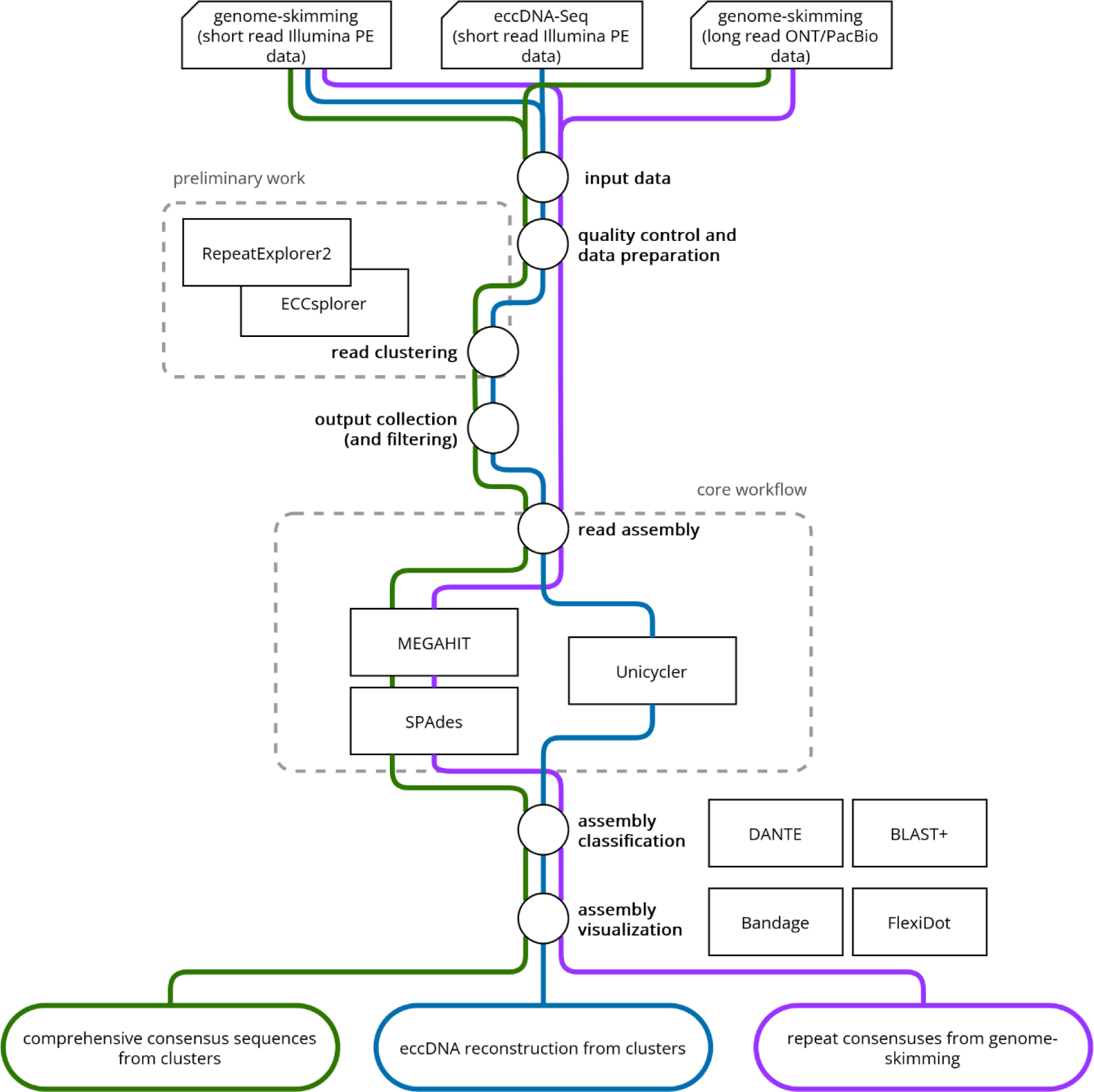
Overview of the repeat assembly workflow. Guided workflows for the creation of comprehensive consensus sequences from repeat clusters (green), the reconstruction of circular sequences from eccDNA candidate clusters from whole genome amplification methods (blue) and the assembly of consensus sequences from highly abundant repeats to explore structural features to solve specific repeat-derived questions (purple).

### Create comprehensive consensus sequences from repeat clusters

This part of the repeat assembly workflow (Fig. 1, green) uses genome-skimming data (short or long reads, example data: short reads from *Beta corolliflora*, 0.1× genome representation) that already has been clustered by RE2 [4, 12] to create comprehensive consensus sequences using a combination of MEGAHIT [19] and SPAdes [20]. After quality control and trimming, the data was processed according to the RE2 protocol using the web-based (galaxy) version or a local installation (--keep-names enabled). Manual correction of the annotation was performed. Reads from the superclusters were collected as fasta, fastq, or as ‘contigs-based’ fastq reads in the following way: Reads in fasta format were used from superclusters as is. Reads in fastq format have been collected from the raw reads using the supercluster reads as reads list. Contig-based reads have been collected from raw read data using only reads that mapped against the supercluster contigs with bowtie2 [21]. Each supercluster read set in each format has been assembled using MEGAHIT (meta-large preset) in a first assembly round. The second assembly round was done with SPAdes (--isolate, --cov-cutoff auto) using the final MEGAHIT contigs and the supercluster contigs as trusted contigs. For a more detailed and guided workflow and access to custom python scripts see the GitHub repository (https://github.com/crimBubble/repeats_and_circles_assembly).

For the example data of *Beta corolliflora,* the statistical data of assembled sequences was collected with seqkit [22] and visualized with ggplot2 [23]. Assembled sequences from reads in fasta format were compared graphically using FlexiDot [24] and sequence-wise using blast+ [25] with the original RE2 contigs, ONT long reads (representing real repeat sequences) and a repeat data base (see section Availability of data). Protein domains have been annotated with DANTE [26] according to the domains specified in the REXdb [27]. Additionally, a score has been calculated for each consensus sequence, defined as length (bp) multiplied by coverage (as reported by SPAdes or RE2). For each repetitive element, the ten consensus sequences with the highest scores were considered the most representative ones and have been selected for comparative statistics and visualizations.

### Reconstruct circular sequences from eccDNA candidate clusters

The second part of the repeat assembly workflow (Fig. 1, blue) reconstructs complete circular sequences following the ECCsplorer pipline [13] with Unicycler [28]. For this, eccDNA-Seq data is used as input (short read, example data: short reads from *Beta vulgaris*, circle-enriched and control data). After quality control and trimming, data were processed according to the ECCsplorer protocol. Manual correction of the annotation and filtering was performed (note that filtering of mitochondrial clusters was omitted compared to the detailed instructions). From the eccDNA candidate superclusters, reads were collected as fastq reads and as contigs-based fastq reads in two ways: Reads in fastq format have been collected from the raw reads using the supercluster reads as reads list. Contig-based reads have been collected from raw reads, using only those that mapped against the supercluster contigs with bowtie2 [21]. For each eccDNA candidate supercluster, the reads in both formats and a combination of both read sets have been assembled with Unicycler in normal mode (--min_fasta_length 1, -- keep 2). Additionally, circles were detected with a custom python script using the networx package. For a more detailed and guided workflow and access to custom python scripts see the GitHub repository (https://github.com/crimBubble/repeats_and_circles_assembly).

For the example data of *Beta vulgaris,* the assembled sequences were compared graphically using FlexiDot [24] and the Bandage assembly viewer [29] with the NCBI reference sequences of the *Beta vulgaris* mitochondrial mini-circles a, d, pO.

### Assemble consensus sequences from highly abundant repeats and explore structural features

The last part of the repeat assembly workflow (Fig. 1, purple) aims at exploring specific questions regarding repeat organization and structure, such understanding linkage or separation of certain repetitive DNAs. Here, we combine MEGAHIT [19] and SPAdes [20] to directly run on genome-skimming reads (short or long reads possible; example data: short reads from *Artemisia annua* and *Tragopogon porrifolius*). After quality control and trimming, data were directly assembled with MEGAHIT (meta-large preset) in a first assembly round. A second assembly round was done with SPAdes (--isolate, --cov-cutoff 20) using the final MEGAHIT contigs as trusted contigs. For a more detailed and guided workflow and access to custom python scripts see the GitHub repository (https://github.com/crimBubble/repeats_and_circles_assembly).

## Results and discussion

### Use case 1: Repeat assembly from *Beta corolliflora* superclusters reveals improved continuity of consensus sequences representing real repetitive elements

The first use case in this study represents the capability of the described workflow to create high-confidence and comprehensive consensus sequences from repetitive elements. Here, we use available data from the recently sequenced *Beta corolliflora,* a wild beet species and close relative of the crop plant sugar beet (*Beta vulgaris*). The consensus sequences have been assembled after clustering of the read data with RE2 [12].

There are multiple ways to retrieve reads from RE2 superclusters for use in our workflow (also see Methods and Implementation). These three ways extract different levels of read information, as follows: (i) “fasta”: read information only; (ii) “fastq”: read and quality information; (iii) “contig-based”: read and quality information based on RE2-contigs from original data (non-sub-sampled data). All three ways have been tested and compared against each other (Fig. 2a; with the colour gradient from orange to green representing repeat abundance). Overall, reads from superclusters in fasta and fastq format performed very similarly, with the fastq reads showing a slight advantage for large superclusters (high read count; Fig. 2a, shades of orange). The contig-based reads showed a higher number of sequences (count of contigs per supercluster) along with the related low N50 values. However, for smaller superclusters (low read count; Fig. 2a, shades of green), the contig-based reads resulted in better consensuses as the standard reads sometimes failed to produce consensus sequences at all. In some cases, SPAdes failed to produce appropriate (at least 100 bp long) consensus sequences (NODEs, e.g. Fig. 2b SCL012). In such cases, the MEGAHIT output usually contained consensuses that instead may be used for downstream analysis (not shown). Taken together, depending on the abundance and characteristics of the repeats of interest, either the supercluster reads in fastq or the contig-based reads show the most useful results. A combination of both read sets can yield additional information (see case study 2). Hence, we conclude that while read input does strongly influence the results, the appropriate choice of reads can be different for individual repetitive elements. As our workflow can be heavily parallelized, all setups can be tested in short time providing the optimal results for each genomic element. In the following, for evaluation of the results, we use read input option (i) “fasta”.

**Fig. 2:**
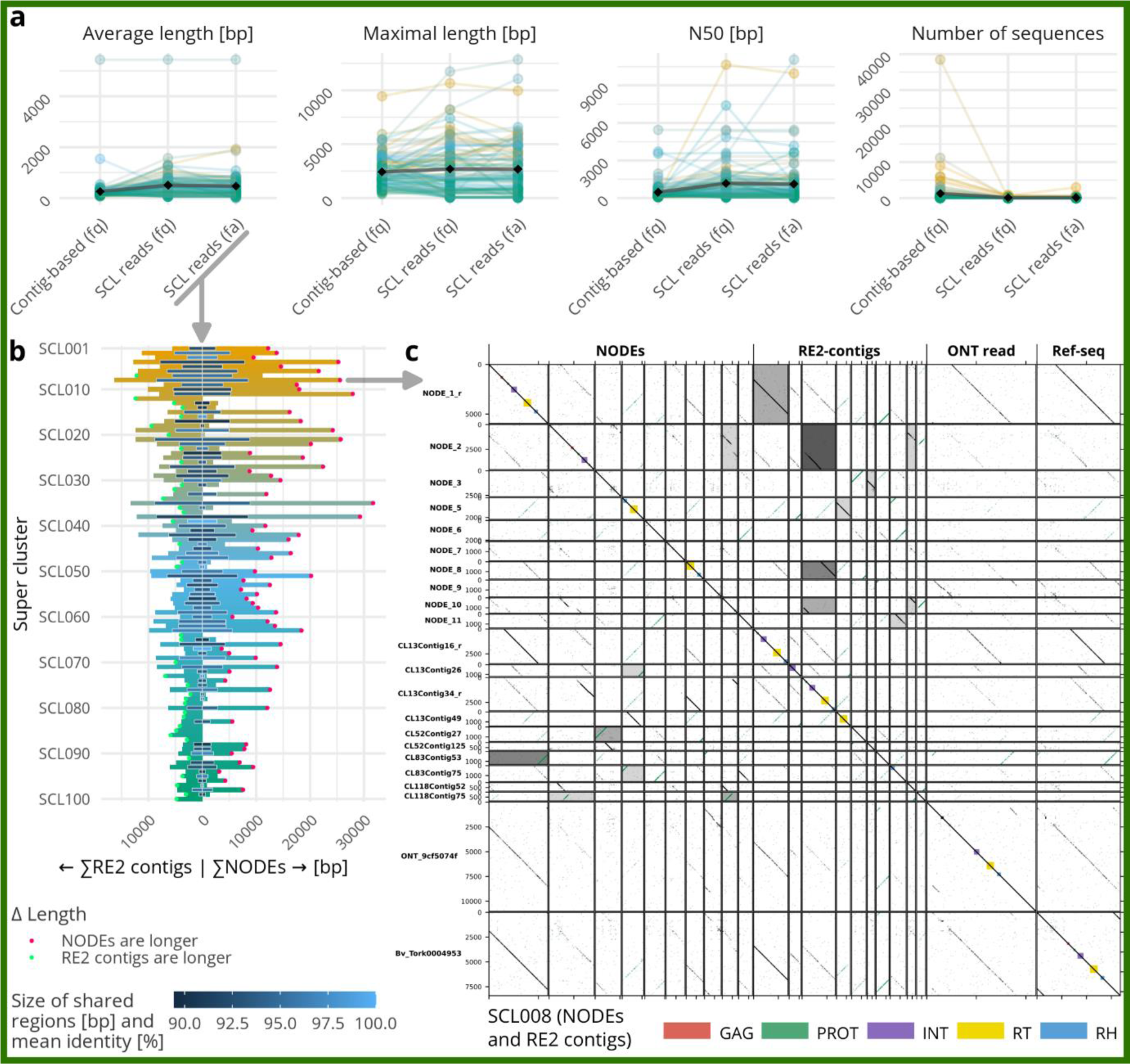
Automated repeat assembly from RepeatExplorer2 (RE2)-derived superclusters (SCLs) leads to long, continuous and accurate repeat consensus sequences.

To assess overall performance of our assembly workflow, we compare the newly generated consensus sequences, in the following referred to as NODEs, with the corresponding RE2-contigs. The Top Ten best-scoring NODEs were overall longer than the corresponding RE2-contigs. Only if NODE generation failed, RE2-contigs were longer (Fig. 2b, side-by-side comparison). Usually, both showed sequence overlaps with a variable length and high sequence similarity (0.4–17.7 kb; 88–100 % identity; see Fig. 2b, length and shading of the central blue bar). For most superclusters, at least one long NODE was produced that was significantly longer than the longest comparable RE2-contig, hence, resulting in a higher continuity of the consensus sequence. Comparing all of the 100 most abundant repetitive DNAs, represented by the Top 100 superclusters, we find that 63 produced more continuous NODEs (Fig. 2b, left, indicated with a red dot) as opposed to 37 more continuous RE2-contigs (Fig. 2b, right, indicated with a green dot).

Exemplarily, we illustrate both, NODEs and RE2-contigs for supercluster 8 (SCL8), representing an Angela-type LTR retrotransposon family (Fig. 2c). In the dotplots, the high fragmentation of the RE2-contigs becomes visible, whereas, for the NODEs, only two sequences are needed to represent the complete retrotransposon (NODE_1_r and NODE_9, Fig. 2c). To further verify the accuracy of the NODEs, we compared them to real repeat representatives (derived from ONT long reads) and to consensus sequences from a *Beta vulgaris* repeat library. In the representative example of SCL8, a real Angela (Tork)-type retrotransposon is highly similar to the calculated NODE sequence (Fig. 2c).

Investigating the architecture of the NODEs more closely, we find that the longest NODEs mostly represent the more conserved parts of a repetitive element, which are usually the protein-coding domains in the case of transposable elements, and the genes in case of rDNAs. The long terminal repeats (LTRs) of the most abundant repetitive elements in plant genomes are less conserved over single retrotransposon families and are present in NODEs with lower scores (i.e., higher NODE numbers, such as NODE_9 of SCL008, Fig. 2c). However, even these less conserved parts are assembled using the presented workflow and usually can be visualised in the bandage graph representations as regions with more branching.

Overall, the assembled NODEs can serve as an additional layer of information when analysing the repetitive DNA of non-model organisms and can be used as a database for transposable elements in following genome assemblies or in the analysis of closely related species. The consensus sequences are especially useful for identifying and classifying individual copies of repetitive elements in subsequent analyses. The consensus sequences can serve as a repeat database reducing the amount of manual work by a) providing more continuous sequences and by b) reducing the amount of sequences which need manual curation. Moreover, a deeper knowledge on the repetitive fraction of a genome might also help to design whole genome assembly strategies: Since this fraction is a major hitch in most assembly approaches, educated decisions on suited technologies and appropriate sequencing coverages are profound and can be addressed with the presented workflow. There is a huge difference in needed data for a genome with, e.g., many conserved sequences in tandem versus a genome with, e.g., many dispersed and less conserved transposon sequences. By providing this information, the RE2 combined with assembly tools can be used to make an optimal use of sequencing budgets by leveraging the different advantages of the latest sequencing technologies such as whale-long ONT reads, high-fidelity PacBio reads, or structural advanced Hi-C conformation capturing.

**(a)** Impact of read input: Comparison of repeat assembly quality measures, when using different input reads for the MEGAHIT/SPAdes assembly workflow. There are three different ways to collect input reads for the assembly workflow: (i) Reads in fasta format can be directly used from the superclusters (SCL reads; fa); (ii) reads in fastq format can be selected from the original read data using the reads listed in the supercluster (SCL reads; fq); (iii) or the reads in fastq format can be selected from the original data based on their similarity to the supercluster contigs (contig-based; fq). Each colour represents a supercluster, displaying the 100 largest superclusters, with colour being on a continuous scale as outlined in Fig. 2b. Black diamonds show mean values.
**(b)** Impact of the new MEGAHIT/SPAdes workflow on consensus length: For the most abundant 100 superclusters (SCLs), we compared the combined length of the assembled consensuses (NODEs; right-facing bars) and RE2-contigs (left-facing bars) using the ten best-scoring contigs/NODEs of each supercluster. The central blue bar indicates the length of the shared sequences between both, NODEs and RE2-contigs, whereas the depth of the blue shade indicates their mean sequence identity. If the NODE assembly produces longer consensuses, this was indicated by a blue dot, whereas a superior RE2 assembly was marked by a green dot.
**(c)** The accuracy of the generated consensus is illustrated by an in-depth view into a selected repeat family, an Angela-type retrotransposon, represented by supercluster 8 (SCL8): Dotplot comparison of the 10 highest-scored NODEs and RE2-contigs from supercluster 8 (SCL008), as well as an ONT long read with an actual Angela copy and a reference sequence for a more detailed sequence-wise comparison. The shading refers to the longest common subsequence (LCS), in which darker grey indicates a longer sequence overlap. In the upper part (above the main diagonal) the forward LCS and in the lower part (below the main diagonal) the reverse LCS is used for shading. The ending “_r” indicates sequences that are displayed as reverse complement.

### Use case 2: Automated reconstruction of mitochondrial mini-circles from *Beta vulgaris* **following the analysis with the ECCsplorer pipeline**

The ECCsplorer pipeline is a tool for the detection of eccDNA that is useful for non-model-organisms as it does not rely on a reference genome assembly. Instead, two short read datasets are compared – one is enriched in eccDNAs, whereas the other is a canonical whole genome shotgun read set. To determine the enriched eccDNA candidates, the ECCsplorer uses RE2 for clustering, similar to use case 1. Hence, it mostly lacks the discovery of complete circular sequences that represent closed DNA rings. Using the presented repeat assembly workflow in addition to the ECCsplorer pipeline solves this problem. To demonstrate this, we use an exemplary dataset with well-characterized eccDNAs (Mann et al. 2022). This dataset is from sugar beet (*Beta vulgaris*) and is enriched in so-called mitochondrial mini-circles (Mann et al. 2022). There are three beet mini-circles in total, named a, d, and p0 that contain shared regions [30, 31]. Running the ECCsplorer on this sample dataset yields a single supercluster with sequences from all three mini-circles, though reconstruction of full circular sequences is failing. Now, with the new MEGAHIT-SPAdes assembly workflow, we could fully retrieve all three complete circular sequences from just this one single supercluster (Fig. 3). The assembled circles are almost identical to the references, sequenced in the 1980s [30, 31], only varying by a few SNPs. These SNPs are likely biological differences rather than actual assembly errors as the mitochondrial DNAs typically have higher mutation rates than genomic DNAs.

**Fig. 3:**
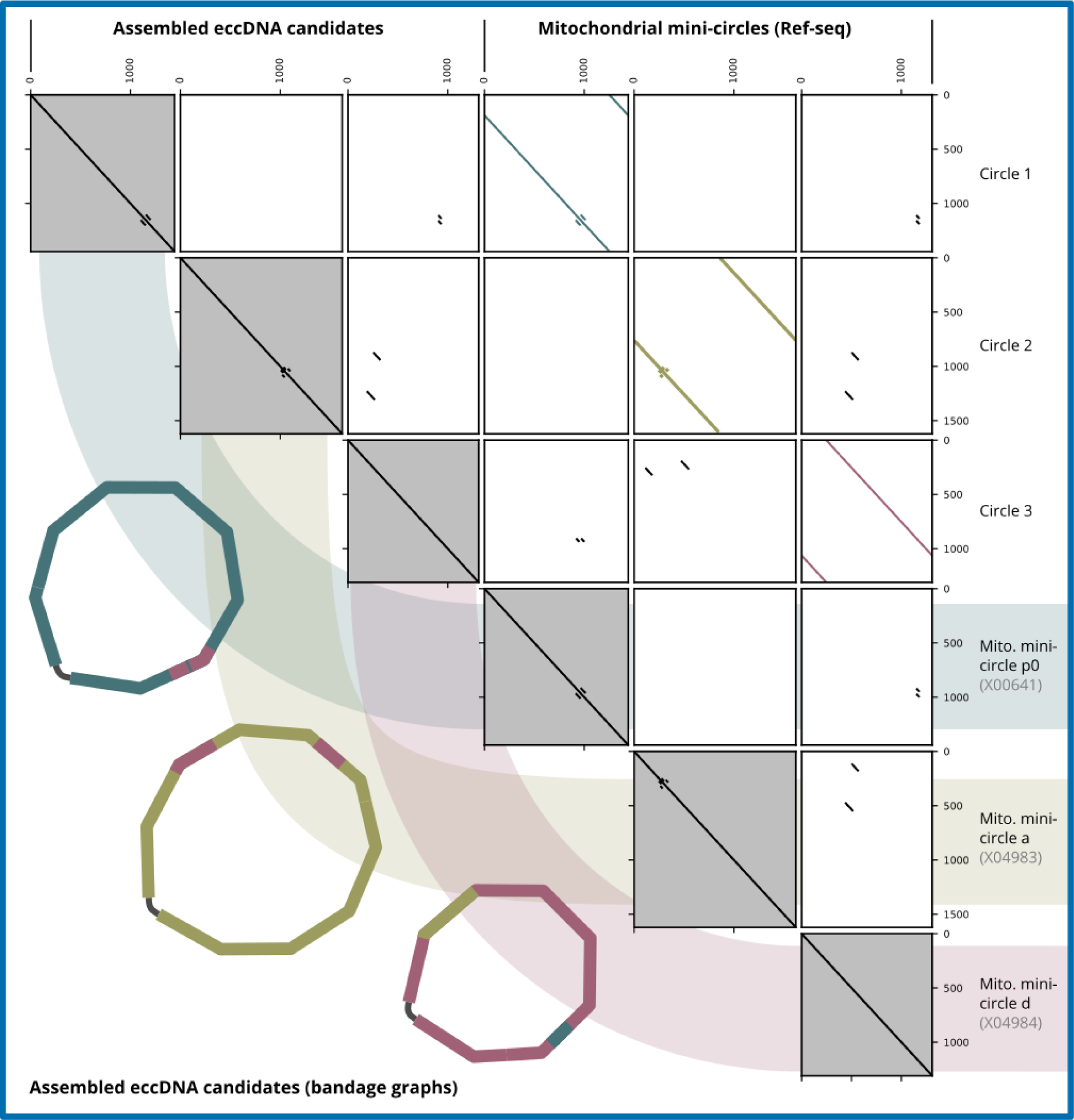
Mitochondrial mini-circles assembled using ECCsplorer (clustering module) output. The dotplots show very high similarity to the published reference sequences and the bandage graph representations show that the full circular sequences could be assembled in high accuracy.

Overall, this shows that the presented workflow is capable of discovering complete circular sequences from eccDNA-Seq data that has been clustered with the ECCsplorer (and RE2) without the need of any reference sequence. This will be especially helpful for the analysis of chimeric eccDNAs, which have been reported recently [32, 33]. Further, we believe that also the bandage graphs and coverage information produced during the workflow can assist in the understanding and differentiation of real chimeric eccDNAs versus template switches of the phi29 polymerase [34–36], without the immediate need for long read sequencing. Template switches might show similar clustering results as the overlapping mini-circles, but our workflow was still able to differentiate between all three individual circular sequences.

Therefore, we predict that using the ECCsplorer followed by the presented workflow is an improvement over current eccDNA assembly methods.

### Use case 3: Exploring the linkage and separation of rDNA in Asteraceae species

The repeat assembly workflow introduced here, may also be helpful in resolving unusual repeat organization patterns. For example, among the Asteraceae, some species harbour an unusual linkage of the 5S and 35S rDNAs [18]. This linkage is often confirmed by fluorescence *in situ* hybridisation (FISH), a powerful, but time-consuming and complex method, relying on specialized equipment and trained staff. With the recent oversupply of sequencing data, the question of rDNA linkage or separation can also be answered using our workflow. The repeat assembly workflow represents a quick and reliable method to confirm linkage or separation of rDNA including the possibility to create consensus sequences. Here, we used data from two Asteraceae species with one of them showing a known linkage of rDNA (*Artemisia annua*, L-type) and the other showing a separation of 5S and 35S rDNA (*Tragopogon porrifolius*, S-type). From the resulting bandage graphs the linkage in *Artemisia annua* and separation in *Tragopogon porrifolius* is clearly visible (Fig. 4). Additionally, the rDNA genes are highly conserved, whereas the spacer regions show some higher variability, indicated by branching. The variability in *Tragopogon porrifolius* is even higher and it might be possible that there are some TE insertions in some 35S rDNA copies which would not be unusual. Furthermore, there is also some variation in the rDNA genes of *Tragopogon porrifolius* that may reflect the additional rDNA loci of both 35S and 5S rDNA [37]. Of course, many derived questions that target homogenized repeat co-occurrences can be targeted as well by the proposed repeat assembly and visualization algorithm (e.g. transposable elements in rDNA, or similar).

**Fig. 4:**
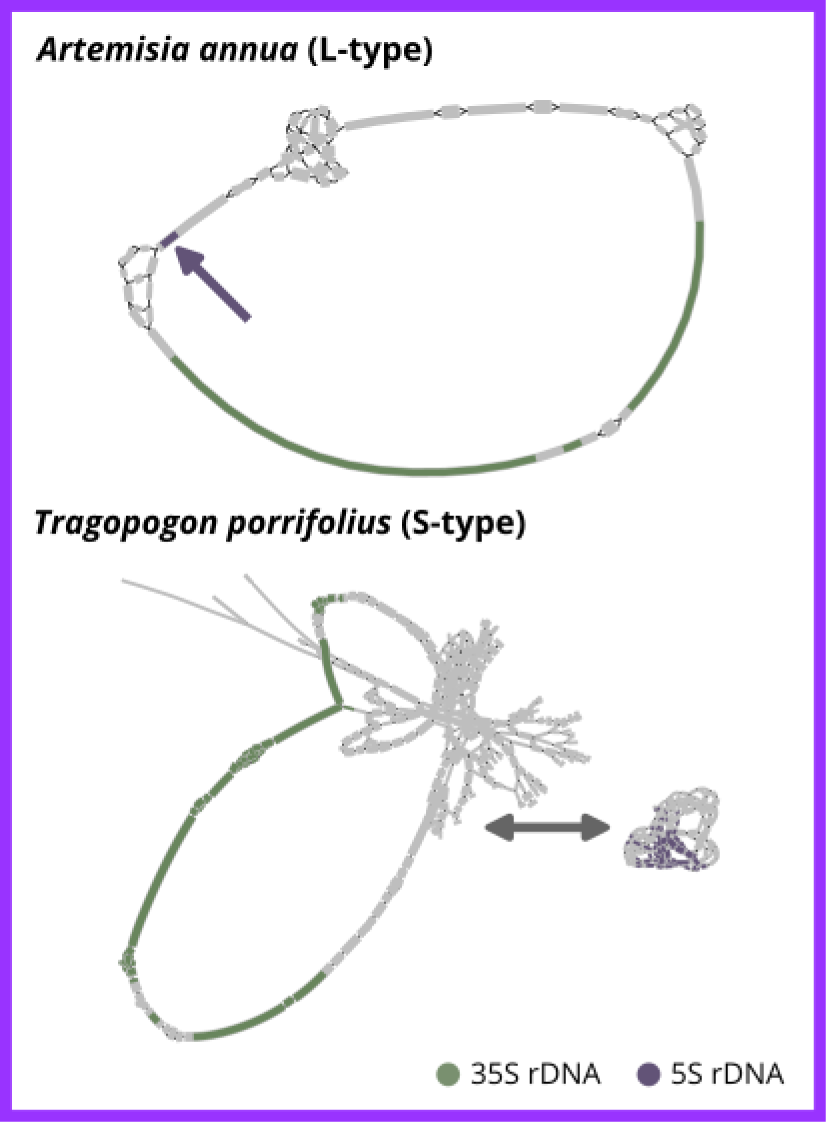
Graphical representation of the repeat consensus assembly (bandage graphs) from genome-skimming data confirms the linkage or separation of rDNA in *Artemisia annua* and *Tragopogon porrifolius*.

## Advantages and limitations

Overall, the present workflows build on existing and well-established methods and extend them in a reproducible manner to provide guidance for follow-up analyses. Manual workloads are reduced wherever possible, allowing automation of the time-consuming manual repeat curation processes.

The new repeat assembly building on RE2 supercluster information assists and automates the manual work that is usually carried out after an RE2 run. Our workflow provides a consistent methodology and is less dependent on previous knowledge about the analysis. Therefore, the presented workflow represents an easy and time-saving way to gain more information from existing data. However, even with these advances, the presented workflow may not always provide finalised, complete consensus sequences, but rather more continuous sequences compared to the original contigs. To reach the best consensus sequences, it may be necessary to try different read collection methods as instructed in use case 1 (Fig. 2a).

The reconstruction of circular sequences from the clustering workflow provides the only described method for the *de novo* detection of eccDNA sequences from short read eccDNA-Seq data so far. Nevertheless, if multiple circles cluster together, the reconstruction still can be challenging and might need an experienced user. However, this is, as demonstrated, still possible and opens up new possibilities for the eccDNA detection and classification.

Exploring structural, chromosomal or chromatin-related features of highly abundant repeats can be quite challenging and often needs time- and cost-intensive experimental methods such as FISH. Recently Garcia et al. (2023) showed how structural repeat features can also be detected by RE2, however, using the presented workflow speeds up the process. Bypassing the RE2 approach completely means that the automated repeat classification is missing, however, allows to input more read data up to 1× coverage without any issues, and thus allows to generate even the most complex rDNA consensus sequences [39]. Also, the use of long reads is possible without any limitations compared to the RE2 approach.

## Conclusion

In this study, three related workflows are presented to assemble repeats and DNA circles from genome skimming and enrichment strategies, such eccDNA-Seq.

Repeats remain one of the major challenges in genome assembly, despite their analysis harboring great potential for understanding genome organization, regulation and evolution. With the presented workflow, we offer a method to generate robust and useful repeat consensuses without extensive manual work. Furthermore, it complements the already existing methods for repeat identification, classification and annotation such as RepeatMasker, LTR_finder, or REPET, which often focus on analyzing already assembled reference genome sequences [40–42]. We conclude that our repeat assembly approach can add large value to read clustering methods that usually only provide an assortment of shorter contigs (use cases 1 and 2). Additionally, repeat assembly without read clustering (use case 3) serves as a faster alternative to answering specific repeat-related questions and to giving insights into structural features. We predict that this workflow will be even more reliable with the upcoming highly accurate sequencing techniques such as PacBio’s Onso short read system.

Extending the usefulness of our repeat assembly approach, we also test it for the *de novo* detection of complete DNA circles. This is useful to understand chimeric eccDNAs and to analyse eccDNAs in non-model organisms. As for the other use cases, the presented workflow builds on existing *de novo* identification and provides examples for advanced uses of the ECCsplorer pipeline. Hence, this approach will generate new insights by providing complete circular consensus sequences for eccDNA candidates.

Overall, the three presented approaches are useful to automate workloads in repeat identification, characterization and annotation, and to overcome the recent data deluge.

## Availability and requirements

Project name: Repeats and circles assembly workflow Project home page: https://github.com/crimBubble/repeats_and_circles_assembly Operating system(s): Linux (tested on Ubuntu 20.04 LTS and 22.04 LTS) Programming language: Shell, Python Other requirements: see GitHub repository License: GLP-3.0

## List of abbreviations

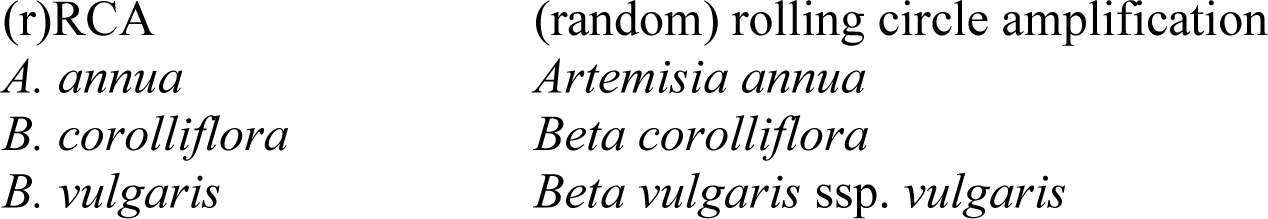
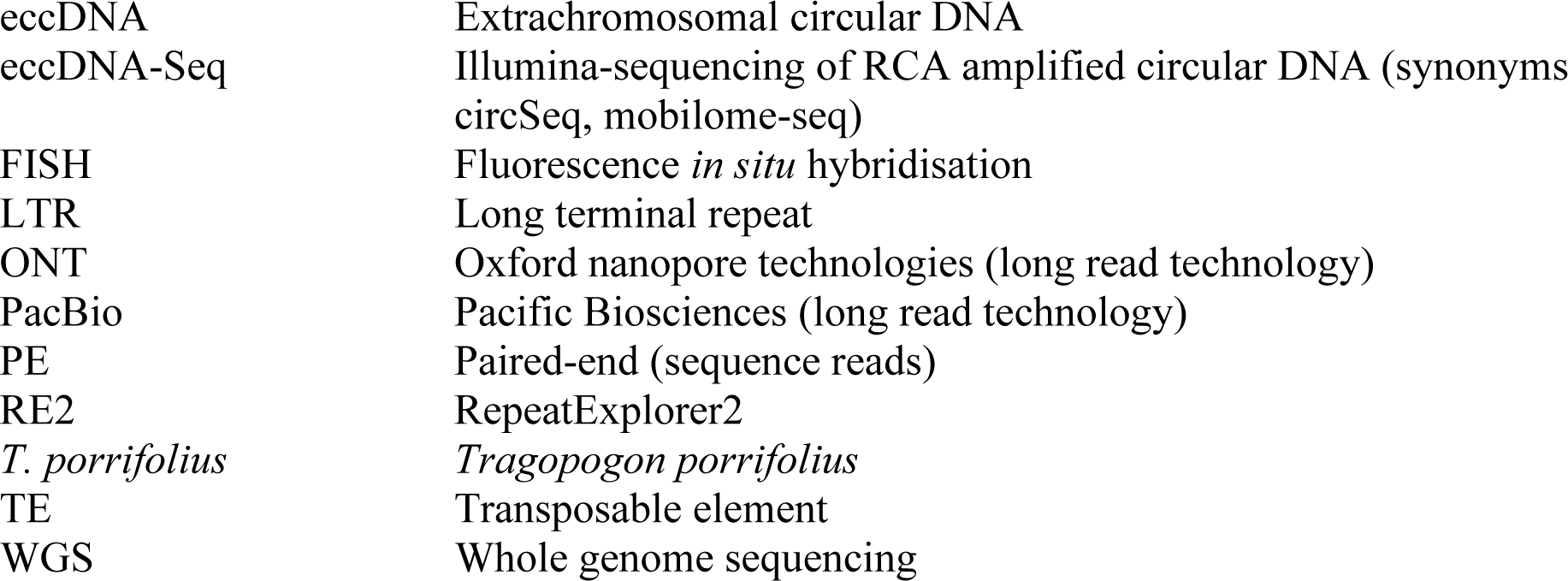

## Declarations

Ethics approval and consent to participate Not applicable.

## Consent for publication

All authors have read and approved this manuscript.

## Availability of data and materials

The datasets together with the accession codes are as follows: *Beta corolliflora* (WGS, Illumina short reads): ERR6110441 (PRJEB45680) [43], the according RepeatExplorer2 run is available at: doi: 10.5281/zenodo.7821055 [4]; *Beta corolliflora* (WGS, ONT long reads): ERR10684093 – ERR10684133 (PRJEB56520) [44]; *Beta vulgaris* (repeat data base): 10.5281/zenodo.8255813; *Beta vulgaris* (eccDNA-Seq, Illumina short reads): ERR6004146 (PRJEB45524) [13]; *Beta vulgaris* (WGS, Illumina short reads): SRR869631 (PRJNA41497) [45]; *Artemisia annua* & *Tragopogon porrifolius* (WGS, Illumina short reads): ERR11535563 and ERR11535566 (PRJEB63080) [39].

## Authors’ contribution

LM wrote the manuscript. LM and KB developed the workflows and wrote the bioinformatic scripts. NS performed the RE2 analysis and provided insights on the repeat landscape. LM and TH conceived the study and designed the experiments. All authors were involved in discussion and editing of the manuscript. All authors read and approved the final manuscript.

## Acknowledgements

We thank the laboratory of B. Weisshaar, and D. Holtgräwe (CeBiTec, Bielefeld) for the publicly available long and short read data used in use case 1. We thank S. Maiwald, S. Garcia, and L. Baader for support and data acquisition of the analyzed *Asteraceae* species. Computational resources were provided by the ELIXIR-CZ project (LM2015047), part of the international ELIXIR infrastructure.

## Figures, tables and additional files

### Supplementary information

Additional file 1: Supplementary Data. Underlying data from Figures 1-3. Contents: Data-Fig1A_B-corolliflora_inputs.tsv, Data-Fig1B_B-corolliflora_contigs.tsv, Data- Fig1C_B-corolliflora-SCL008.fasta, Data-Fig1C_B-corolliflora-SCL008.gff, Data-Fig2_B- vulgaris-mini-circles_assembly.gfa, Data-Fig3_A-annua-rDNA.gfa, Data-Fig3_T-porrifolius- rDNA.gfa.

